# Direct Visualization of Live Zebrafish Glycan via Single-step Metabolic Labeling with Fluorophore-tagged Nucleotide Sugars

**DOI:** 10.1101/548016

**Authors:** Senlian Hong, Pankaj Sahai-Hernandez, Digantkumar Gopaldas Chapla, Kelley W. Moremen, David Traver, Peng Wu

**Author notes:** These authors contributed equally to this work (Senlian Hong, Pankaj Sahai-Hernandez).

## Abstract

Dynamic turnover of cell-surface glycans is involved in a myriad of biological events, making this process an attractive target for in vivo molecular imaging. Metabolic glycan labeling coupled with ‘bioorthogonal chemistry’ has paved the way for visualizing glycans in living organisms. However, a two-step labeling sequence is required, which is prone to tissue penetration difficulties for the imaging probes. Here, by exploring the substrate promiscuity of endogenous glycosyltransferases, we developed a single-step fluorescent glycan labeling strategy by directly using fluorophore-tagged analogs of the nucleotide sugars. Injecting the fluorophore-tagged sialic acid and fucose into the yolk of zebrafish embryos at the one-cell stage enables systematic imaging of sialylation and fucosylation in live zebrafish embryos at distinct developmental stages. From these studies, we obtained insights into the role of sialylated and fucosylated glycans in zebrafish hematopoiesis.

## Introduction

Metabolic oligosaccharide engineering (MOE) coupled with the bioorthogonal chemical reporter strategy has opened an avenue for labeling and visualizing glycans in living organisms.^1–5^ In this approach, the cell’s glycan biosynthetic machinery is exploited to install a biorthogonal chemical group onto cell surface glycans, which is then covalently labeled in a secondary step with a complementary probe.^6^ However, this strategy has a few intrinsic limitations, such as the biocompatibility of a chosen bioorthogonal reaction and poor deep-tissue penetration of the probes.^7^

Chemoenzymatic glycan editing, however, enables the direct incorporation of a fluorescently labeled monosaccharide without the requirement of a second-step covalent reaction. By using recombinant sialyltransferases (STs) and fucosyltransferases (FTs) with broad donor substrate scopes, sialic acid (Sia) and fucose (Fuc) conjugated with fluorescent dyes can be directly transferred to the cell surface from the corresponding nucleotide sugars.^8–11^ Furthermore, recent findings have shown that fluorescently labeled trehalose and *N*-acetylglucosamine can be directly incorporated into the cell wall of *M*. *tuberculosis*^12^ and intracellular *O*-Glc*N*Acylated proteins in cultured mammalian cells^13^, respectively. Inspired by these findings, here, we sought to explore the feasibility of directly incorporating fluorophore-labeled monosaccharides into the cellular glycans of living organisms by exploiting the substrate promiscuity of endogenous glycosyltransferases (Fig. 1a).

**Figure 1.**
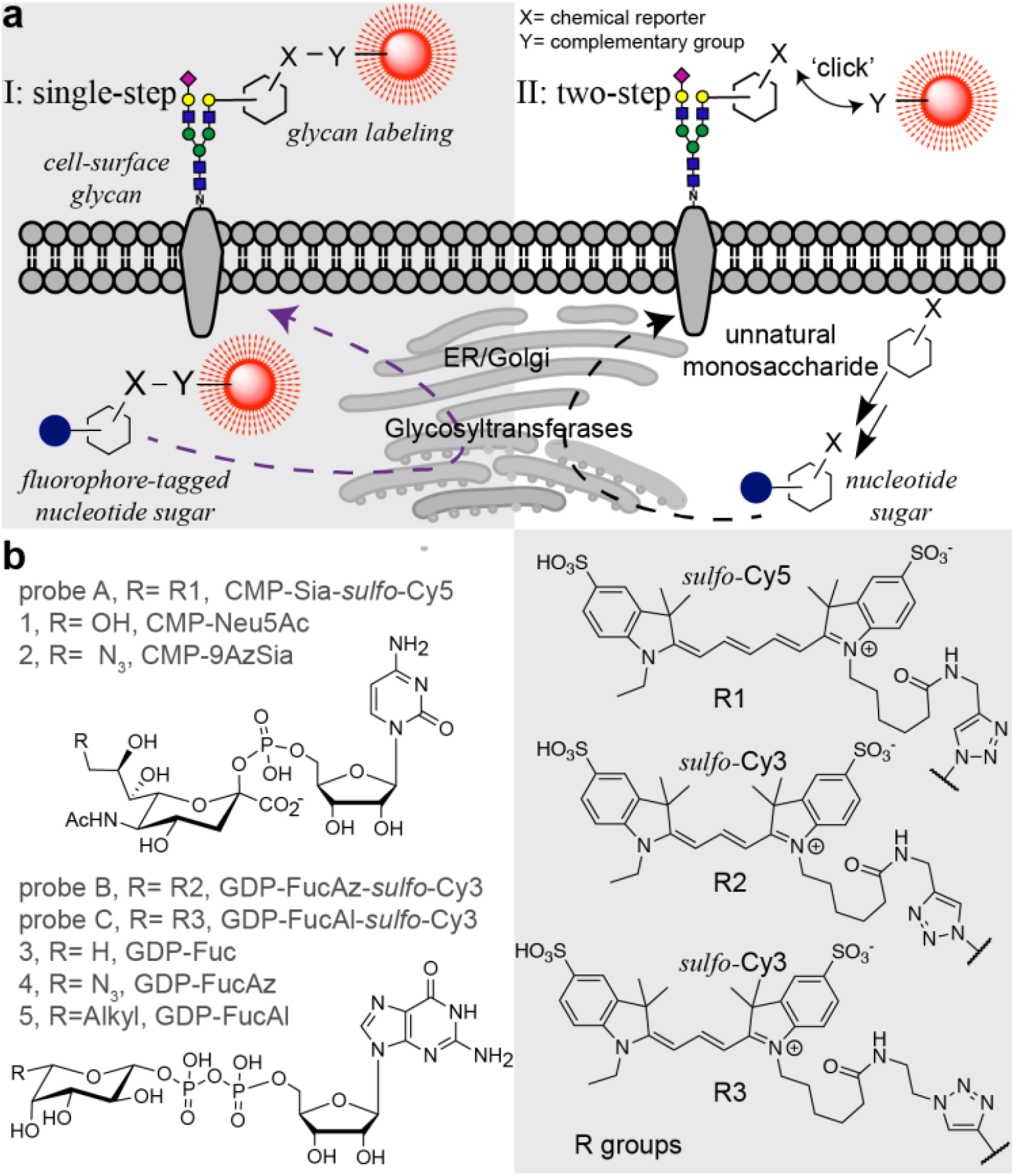
a), Comparison of the single-step fluorescent glycan visualization via metabolic incorporation of unnatural fluorophore-tagged nucleotide sugars versus the conventional, two-step approach. b), Nucleotide sugar analogs functionalized with a chemical reporter (alkyl and azido) or a fluorophore (Cy3 and Cy5) that were used in this study.

## Results and Discussion

We assessed the incorporation of fluorophore-labeled monosaccharides in the live zebrafish embryo due to its optical transparency, external fertilization, and amenability to genetic and embryological manipulations.^14–16^ First, we synthesized a fluorophore-labeled CMP-*N*-acetylneuraminic acid (CMP-Neu5NAc). Previous studies have shown that many sialyltransferases (STs) can tolerate large substituents at C9 of Neu5NAc.^17,18^ Therefore, we synthesized CMP-Sia-*sulfo*-Cy5 (probe A) by coupling CMP-9AzSia^19^ with Al-*sulfo*-Cy5 via ligand (BTTP) assisted copper-catalyzed azide-alkyl [3+2] cycloaddition (CuAAC)^20^.

To first validate that the Cy5-tagged CMP-Sia analog can be incorporated by sialyltransferases in live cells, we examined the feasibility of transferring Sia-*sulfo*-Cy5 onto the cell-surface of sialylation-defect Chinese Hamster Ovary (CHO) mutant Lec2, using three recombinant human STs (hST3Gal1, hST3Gal4, hST6Gal1).^21^ ST6Gal1/2 and ST3Gal 1/2/3/4/5 are evolutionarily conserved in human and zebrafish genomes and STs in both species share high homology,^22–24^ therefore it is reasonable to use the human homologs to examine the donor substrate scope. After incubation with STs and probe A, we detected intense Cy5 fluorescence on Lec2 cells, but not in the cells treated without STs (Fig. s1). Confirming that STs had the capability to incorporate probe A onto cell-surface glycans for the single-step glycan labeling.

Next, we microinjected probe A into the yolk sack of zebrafish embryos at the one-cell stage,^25^ which enables the injected nucleotide sugar to disperse and incorporate into all daughter cells during early zebrafish embryogenesis (Fig. 2a). The injected embryos were subsequently imaged using fluorescent microscope at distinct developmental stages. As shown in the supplemental Fig. s2, overwhelming fluorescent signals in the yolk due to the accumulation of probe A prevented the visualization of the incorporated Sia in other tissue structures during the early embryogenesis. To our delight, the incorporation of probe A in several distinct tissue structures became apparent starting at 14 hours’ post fertilization (hpf), and 28 hpf. Importantly, the probe-dependent Cy5 signal in live embryos is significantly reduced by coinjection of sialylation inhibitor 3FaxNeu5Ac^26^ or natural sialylation precursor CMP-Neu5Ac with probe A (Fig. s3), suggesting specific labeling of cell-surface glycans in zebrafish.

**Figure 2.**
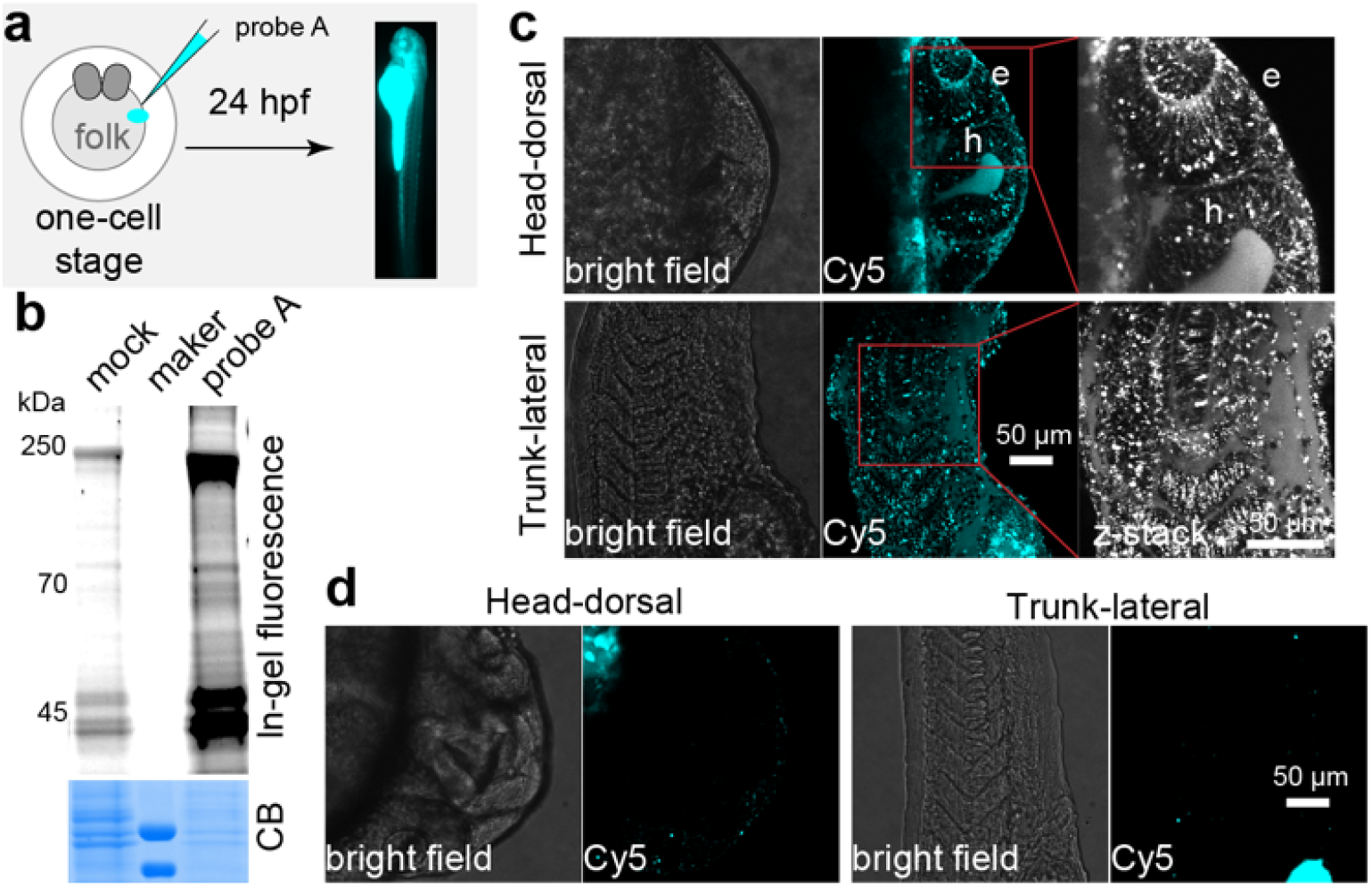
Metabolic incorporation of CMP-Sia-*sulfo*-Cy5 (probe A) onto sialylated glycan in live zebrafish embryos. a), Workflow of using probe A to fluorescently label sialylated glycan in zebrafish embryos. After a 24-hour incubation, the labeled glycoconjugates in live embryos were directly imaged. b), In-gel analysis of Cy5 fluorescence of labeled glycoproteins (top panel). The same counts of embryos were collected, deyolked, and the resulted lysis was loaded onto SDS-PAGE. The loading was evaluated by Coomassie blue staining (CB). c, d), The confocal images of embryos injected with probe A (c), or prequenched reaction mixture (d, mock control). Intensive Cy5 fluorescence was detected in the eye (e) and hindbrain (h) regions of the head, in the somatic tissue, muscle cells, and intro-hypochord of the trunk.

To confirm that probe A was metabolically incorporated into zebrafish glycoproteins, the lysates of the 24 hpf deyolked embryos were analyzed using an in-gel fluorescence assay. We detected strong A-dependent Cy5 signal in SDS-PAGE resolved proteins with a molecular weight ranging from 30-250 kD (Fig. 2b and s4), which was subsequently removed by PNGase F-assisted releasing of N-glycans or neuraminidase-directed hydrolyzing Sia from glycoproteins (Fig. s5). Together, these results strongly suggest that CMP-Sia-*sulfo*-Cy5 is metabolically incorporated in sialylated glycoproteins.

Once we confirmed the specific labeling of probe A into the cell-surface glycoproteins, we proceeded to perform systematic imaging of sialylation in zebrafish tissues at 24 hpf. We observed distinct Cy5 fluorescence in the head and trunk regions (Fig. 2c; movie s1 and s2). Specifically, in the eye and hindbrain regions of the brain, the hypochord in the trunk, and muscle cells in the somites exhibited strong Cy5 fluorescence. Importantly, in the control embryos that were injected with CMP-9AzSia and Al-*sulfo*-Cy5, only negligible background Cy5 signals was detected (Fig. 2d). These observations are consistent with what previously been reported by Bertozzi and coworkers,^17,18^ in which BCNSia was used as the metabolic substrate and the detection was realized by the reaction with a fluorogenic tetrazine probe injected into the posterior caudal vein. In this approach, imaging can only be launched at 30 hpf, which is the earliest time that the caudal vein starts to form. Since the incorporation of unnatural analogs onto glycans could potentially interfere with their biological functions, we further analyzed the morphological defects of zebrafish embryos injected with different concentrations of probe A (2.5, 5 or 10 pmol). However, at 24 hpf we did not observe any apparent morphological defects in any of the injected embryos (Fig. s6), suggesting that probe A do not interfere with normal biological functions.

Due to the toxicity of probe B, systematic imaging of fucosylation was conducted with probe C, even though probe C showed weaker incorporation. Confocal images of live zebrafish embryos injected with probe C at 24 hpf revealed a highly-localized distribution of labeled fucosides in the head (lateral, ventral and dorsal views) and trunk regions (lateral view) (Fig. 3 and s 11; movie s5 and s6). Abundant fucosylation was specifically detected within the eye, olfactory placode, midbrain, hindbrain, hypochord, dorsal fin, blood vessels and the caudal hematopoietic tissue (cht). The incorporation appeared more intense in neural structures such as the midbrain and the hindbrain. Interestingly, the midbrain-hindbrain boundary (mhb) did not show any incorporation of the unnatural fucose. Moreover, this strategy also enabled time-lapse tracking of fucosylated glycans in live zebrafish embryos. For example, Cy3-labeled fucoside was found to be abundantly expressed in the retina and the optical cup of eyes, and dynamically moving along the ganglion cells of acetylated microtubules, which are regarded as stable, long-lived microtubules (Fig. 3b 4-t1 and 4-t2; movie s7).

**Figure 3.**
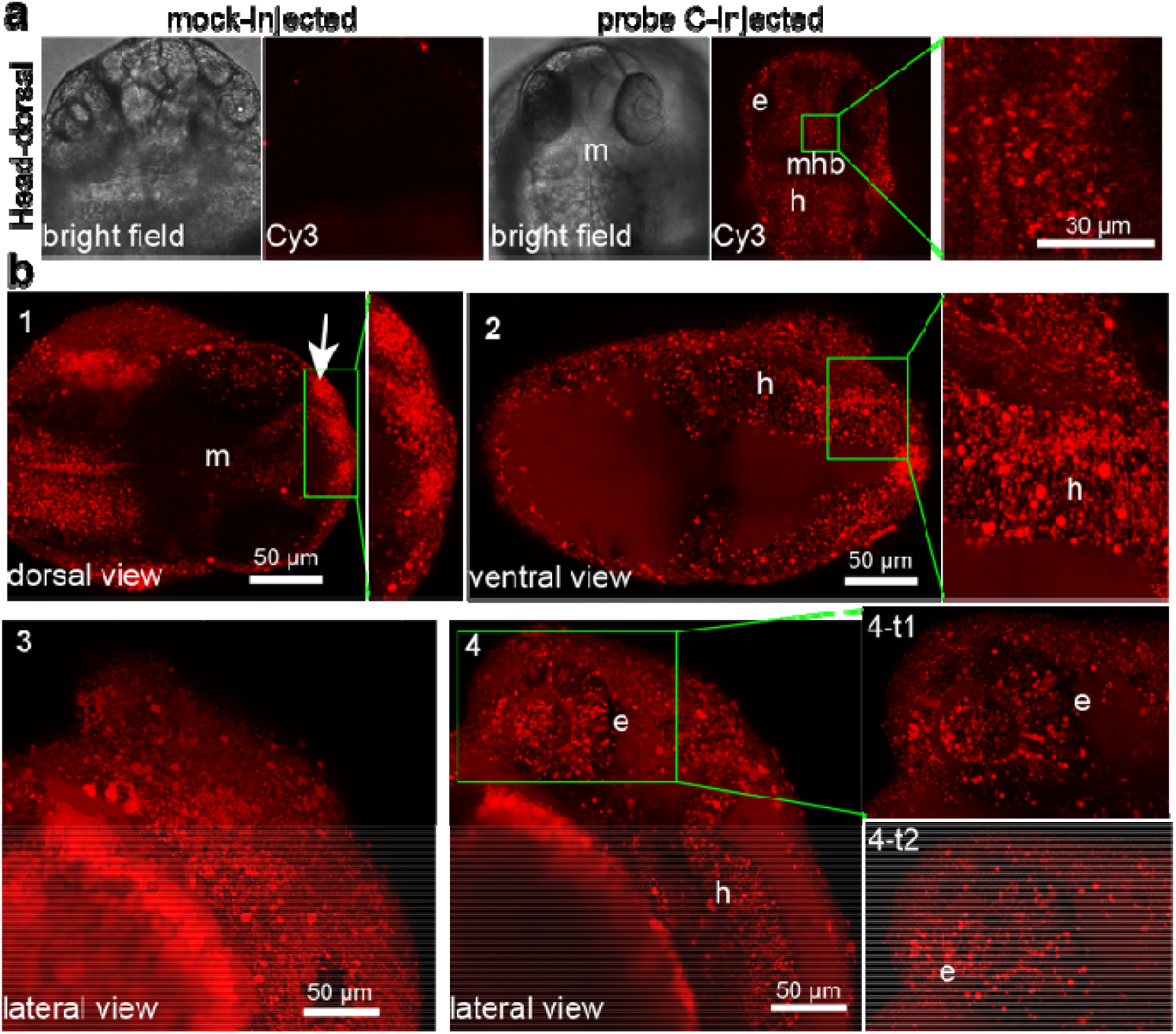
Systematic visualization of fucosylation in live zebrafish by metabolic incorporation of GDP-FucAl-*sulfo*-Cy3 (probe C) onto fucosylated glycans. a), confocal images of embryos injected with probe C or prequenched reaction mixture (mock control). In probe C-injected embryos, strong Cy3 fluorescence was detected in the eye (e), midbrain (m) and hindbrain (h) regions, while only negligible background fluorescence was detected in the midbrain-hindbrain boundary (mhb) as shown in the high magnification of the midbrain inset of head-dorsal view. b), Dorsal view (1 and 2) of a head in different depth, the developing mouth showed intensive fucosylation (arrow). Lateral views of head surface (3) and in deep-tissue at different time points (4, 4-t1 and 4-t2).

The ability to simultaneously visualize distinct biomolecules in vivo is of essential importance for understanding normal and disease processes. We then evaluated the feasibility to visualize sialylated and fucosylated glycans in parallel. To this end, we co-injected probe A and probe C at a 1:1 molar ratio into the yolk of zebrafish embryos and conducted confocal imaging at different time points (Fig. 4a and s12). As shown in Fig. 4a, sialylated glycans have broader distributions than fucosylated counterparts at 24 hpf embryos. The blood vessel is the region showing the strongest co-localization of both glycans. However, these two types of glycans also exhibit many distinct distribution patterns. For example, sialylation was abundantly detected in the notochord and myotome region, while fucosylation was generally localized in the vascular network. In addition, the labeled fucoside decayed at a much faster rate than the labeled sialosides; by 72 hpf, the Cy3-associated signal disappeared almost completely in the brain; whereas abundant Cy5 signal was still detectable at this time (Fig. s13).

**Figure 4.**
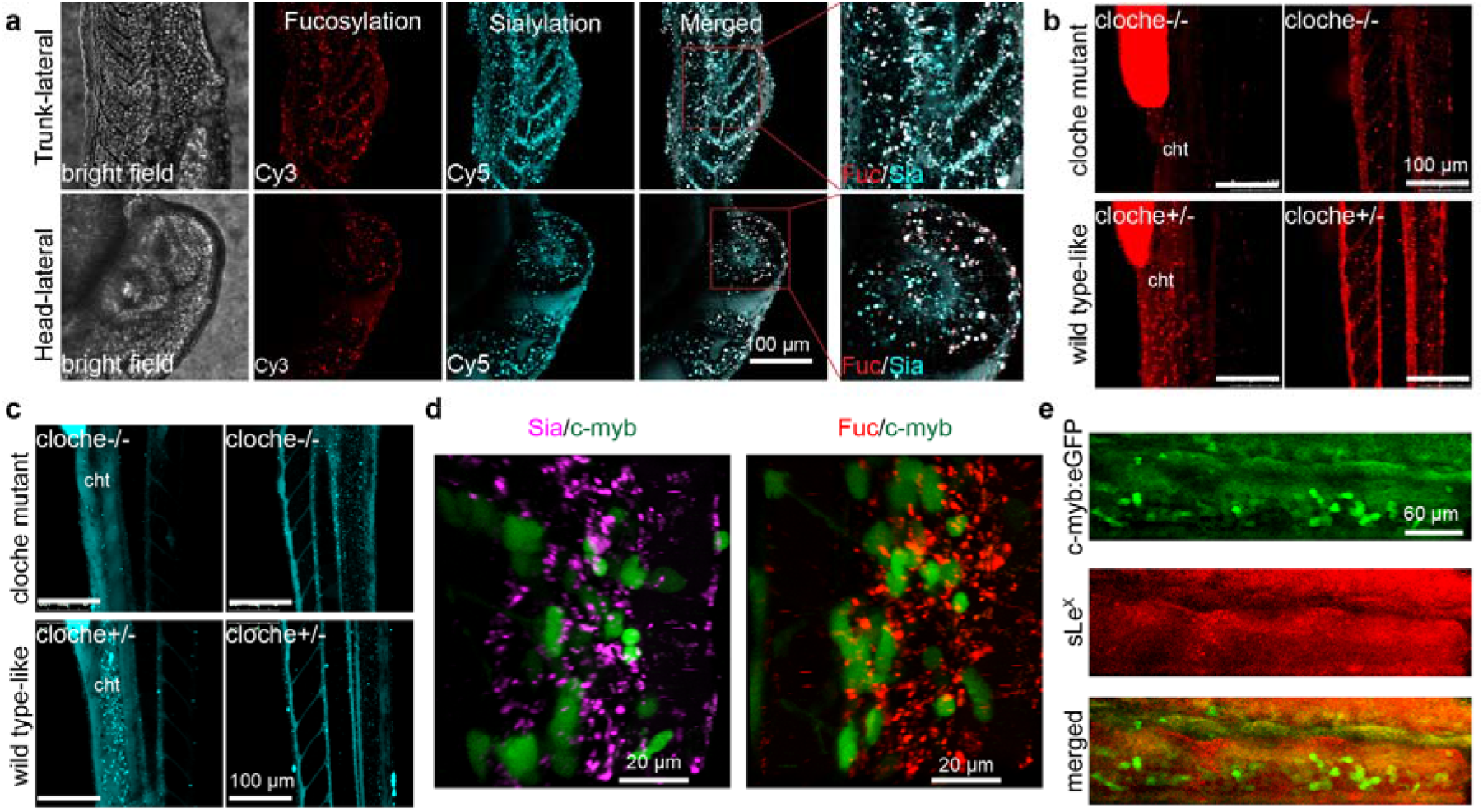
Simultaneous visualization of the deep-tissue fucosylation (Cy3 in red channel) and sialylation (Cy5 in cyan channel). a), The lateral view of zebrafish embryos co-injected with probe A and probe C at 24 hpf. b, c), The trunk (lateral view of cloche mutant (cloche −/−) or wildtype-like siblings (cloche+/−) at 48 hpf was visualized at different depth. The fluorescence in yolk was an endogenic control for the success of microinjections. d), In the cht region of Tg(c-Myb:eGFP) embryos, HSCs (green channel) do not co-localized with the labeled fucosides and sialosides. e), In Tg(c-Myb:eGFP) embryos, HSCs do not co-localize with cells expressing sLe^X^ (red) in the cht region.

Next, we directly compared the labeling patterns of the single-step glycan labeling to the two-step bioorthogonal chemical reporter strategy previously well-established. We co-injected probe C and GDP-FucAz into the same embryo, and the metabolically incorporated FucAz was fluorescently labeled with Al-*sulfo*-Cy5 by BTTPS-assisted CuAAC reaction at 24 hpf. In deep tissues such as the cht region, we only observed the Cy3 signal, but not that of Cy5. Likewise, only a very weak Cy5 signal was detected in the trunk and head regions, which is likely due to the poor permeability of labeling reagents into the deep-tissue of live zebrafish embryos. The observation that a much stronger Cy5 signal was detected at the tip of the tail compared to the Cy3 signal is likely caused by the same issue (Fig. s14). Similar results were also observed in probe C and GDP-FucAl coinjected zebrafish embryos (Fig. s15).

To exclude the possibility that the observed labeling patterns were generated by unincorporated probes circling in the blood or diffusing into the tissue matrix, a hematopoietic mutant line known as ‘Cloche’^27^ was used to repeat our labeling experiments. In this mutant (cloche−/−), no blood circling system including cht, is formed due to the failure to specify all blood cell types in the developing embryos. We injected embryos from cloche mutant heterozygous (cloche+/−) parents with probe A or C. A distinct labeling pattern was detected in the trunk and head regions of wildtype-like sibling embryos (cloche+/−) that has normal hemato-vascular cell types, as we had previously observed. In the cloche−/− embryos that are easily distinguished by the lack of blood flow after 26 hpf, the loss of the labeling was observed in tissues such as cht due to the loss of vascular development (Fig. 4b, c, movie s8-s11). Based on these observations, we reasoned that the background fluorescence generated by the free, unincorporated probes in the bloodstream is negligible.

The cht region is a transient vascular network formed by the ramification of the caudal vein plexus, provides a necessary niche microenvironment for supporting the development of nascent hematopoietic stem cells (HSCs).^28^ The cht is of critical importance for the maturation of these blood cell and immune cell types. To explore the possible biological functions of glycans within this compartment, we imaged probe A and probe C-injected zebrafish in the background of hematopoietic marker Tg(cMyb:eGFP)^29^ whose HSCs produced by definitive hematopoiesis can be tracked by the expression of eGFP (Fig. 4d, movie s12, and s13). Definitive hematopoiesis produces multi-potent blood cell types that give rise to multiple lineages through cellular intermediates, and supports the production of all blood cells throughout adulthood. Within the cht region, we observed that the fluorescently labeled sialosides and fucoside were present in cells that are in direct contact with the eGFP-expressing HSCs, but not within the HSCs themselves. Previous studies have reported that sialyl-Lewis X (sLe^X^, Siaα2-3Galβ1-4(Fucα1-3)-GlcNAc) plays critical roles in mediating leukocyte adhesion and lymphocyte homing,^30,31^ we immunostained sLe^X^ epitopes in Tg(c-Myb:eGFP) embryos with an anti-CLA antibody (Fig. 4d). Despite the high fluorescence background, the intensive staining within the cht region was predominantly found in the extracellular matrix, rather than co-localizing with HSCs. This observation strongly suggests that the interactions mediated by sialylated and fucosylated glycans such as sLe^X^ are involved in the hematovascular cell biology in zebrafish.

During these imaging studies, we made an intriguing observation: compared to their sibling embryos (cloche+/−), the incorporation of GDP-Fuc based Probe C was significantly decreased in the mutant embryos (cloche−/−); whereas the incorporation of CMP-Sia-based probe A was essentially unchanged (Fig. 4b and 4c). Interestingly, compared to the untreated or the probe A injected groups approximately 5-fold more cloche−/− mutant embryos were still alive at 48 hpf in the probe C-microinjected group (Fig. s16a). These observations prompted us to examine the expression of fucoside biosynthesis enzymes in the cloche mutants. We performed RT-qPCR analysis of gene expression of key components of the GDP-Fuc *de novo* biosynthesis and salvage pathways (Fig. s17a), and found that the mRNA levels of fucose kinase (FUK) and GDP-Fuc transporter (SLC35C1) were significantly downregulated in the cloche mutants (cloche−/−) (Fig. s17b). Consistent with those changes, the global fucosylation in cloche−/− embryos was significantly down-regulated compared to that of the wildtype embryos or their siblings (cloche+/−) as assessed by fluorescein-conjugated *Aleuria Aurantia* Lectin (AAL-FITC) staining (AAL is a lectin specific for α1-3- and α1-6-linked fucose) (Fig. 5a). By administering GDP-Fuc into embryos of the cloche mutants (cloche−/−) at the one-cell stage, the global fucosylation level can be largely rescued (Fig. 5b).

**Figure 5.**
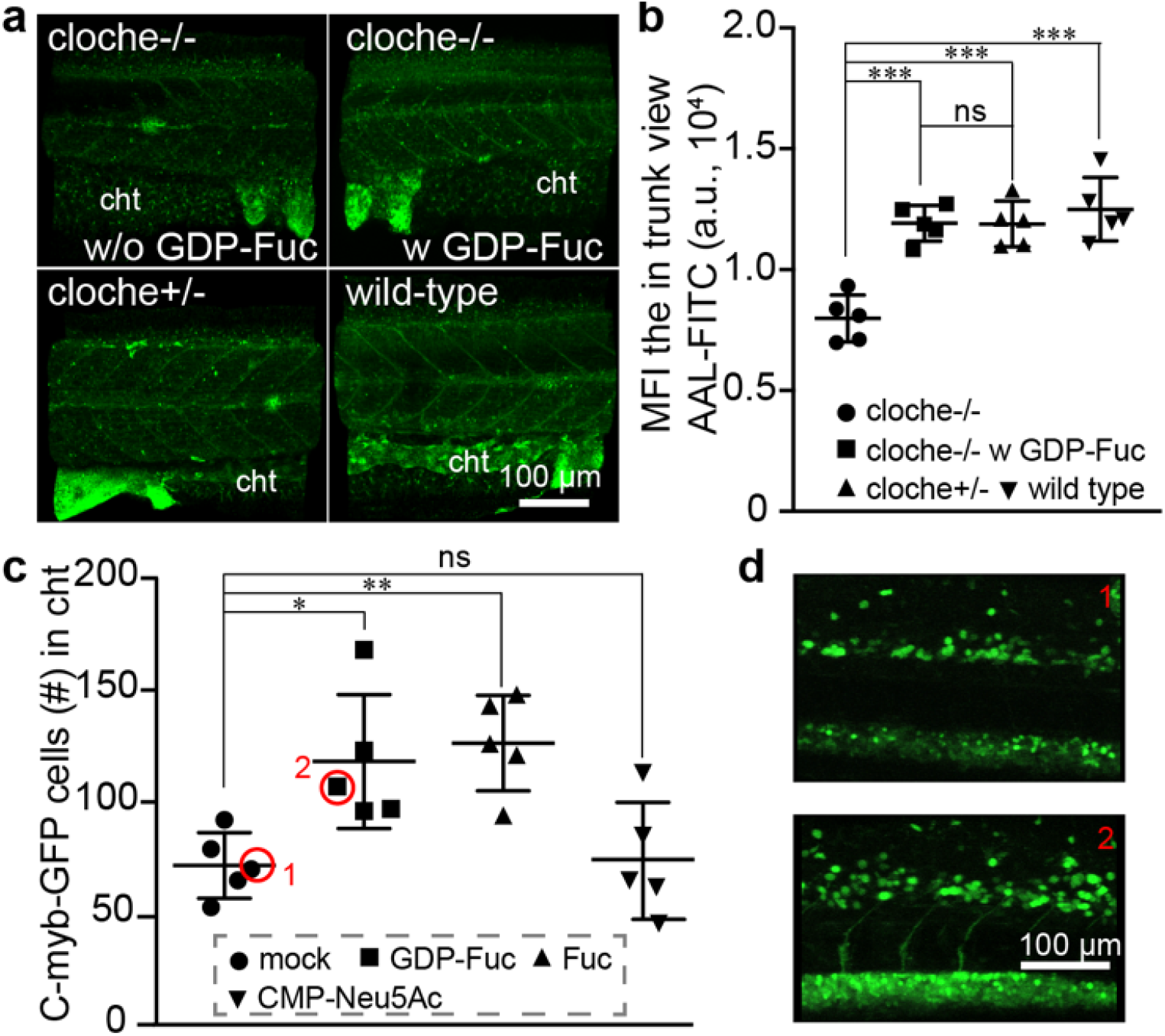
Impairing the global decrease of fucosylation in cloche mutant (cloche−/−) by the administration of the precursor monosaccharide into the yolk of embryos at one-cell stage. a), The 3D confocal images of the fucosylation in the trunk of different embryos that were stained by AAL-FITC. The wild-type, cloche mutant (cloche−/−), wildtype-like sibling (cloche+/−), and GDP-Fuc-injected cloche mutant was collected at 48 hpf and stained with AAL-FITC after fixation and permeabilization. b), The fluorescent intensity in the trunk view of embryos was quantified and presented by mean fluorescence intensity (MFI) via ImageJ. c, d), Confocal imaging-assisted counting of HSCs cells in the cht. Tg(c-Myb:eGFP) embryos at 1-2 cell stage were injected with Fuc, GDP-Fuc, CMP-Neu5Ac, or mock loading buffer, and the HSCs were quantified at 48 hpf. The * represents p<0.05, the ** represents p<0.01, the *** represents p<0.001 and the ns represents not significant in the T-test.

All vertebrates, including zebrafish, has two waves of hematopoiesis^32,33^ The earlier of these is known as the primitive wave and the later one as the definitive wave. The primitive wave is responsible for the production of red blood cell types that can facilitate tissue oxygenation as the embryo undergoes rapid growth; whereas definitive or adult hematopoiesis provides the organism with long-term HSCs that are capable of unlimited self-renewal and generating all mature hematopoietic lineages. The cloche mutant (cloche−/−) exhibits serious defects in both waves, but interestingly the absence of vascular cells is partially rescued by 48 hpf.^27^

To try and decipher the role that fucosylation might play during early zebrafish hematopoietic development, we injected Fuc or GDP-Fuc into transgenic zebrafish lines harboring reporters of these two stages of hematopoiesis and followed their development. Although no convincing findings were made using reporter lines of the primitive wave, in the transgenic zebrafish line Tg(cMyb:eGFP) whose definitive blood progenitors can be directly traced by eGFP-expression, Fuc and GDP-Fuc injection distinctively increased the population of eGFP-labeled HSCs in the cht niche region between 48-72 hpf. These changes were not detectable in mock and CMP-Neu5Ac administrated groups (Fig. 5c and 5d). Finally, we also identified significantly larger number of live cloche−/− mutant embryos 48 hpf after the administration of Fuc or GDP-Fuc compared to CMP-Neu5Ac-treated or mock-treated groups (Fig. s16b).

## Discussion

Previous studies by Bertozzi and us demonstrated that unnatural UDP-GalNAc bearing an azide tag and GDP-fucose bearing an azide or alkyne tag can be used by endogenous ppGalNActransferases and fucosyltransferases, respectively, and incorporated into cellular glycoconjugates in zebrafish embryos^34,35^ Bioorthogonal chemistry can then be employed for their visualization as early as at the two-cell stage. Via this approach, rapid glycan migration to the cleavage furrow of mitotic cells was observed despite poor tissue penetration of fluorescent probes used for their detection.

In the current study, we have demonstrated that fluorophore-tagged CMP-sialic acid and GDP-fucose (probes A-C) injected into the yolk of zebrafish embryos are incorporated onto cellular glycoconjugates, enabling the direct deep-tissue visualization of sialylated and fucosylated glycans. Using this single-step approach, time lapse, high-resolution, and sensitive confocal imaging of these biologically important glycans in zebrafish embryogenesis is realized. Importantly, this approach enables direct visualization of the dynamic turnover of glycans in live embryos. For example, we observed that in the head region, fucosylatd glycans have a quicker turnover rate than sialylated glycans.^24,36,37^

Studies by Stanley, Haltiwanger, and Taniguchi have shown that knockout of FUT8, POFUT1 or POFUT2 is lethal to mice.^38–41^ In addition, congenital mutations of the Golgi localized GDP-fucose transporter SLC35C1 cause leukocyte adhesion deficiency type II, whose manifestation is severe developmental and immune deficiencies.^42^ In line with these previous observations, we discovered that cloche mutation leads to down-regulation of *fuk* and *slc35c1,* and accordingly reduced cell-surface fucosylation, which can be partially compensated by exogenously introduced Fuc or GDP-Fuc. Interestingly, not only did Fuc or GDP-Fuc injection expand the HSC-population in the cht niche, but this treatment also increased the survival of the cloche mutant significantly. Our findings combined with previous reports^43–46^ underscore the critical role of fucosylation in hematopoiesis, especially in the definitive wave, though the exact molecular mechanism for our observed phenotypes is still waiting to be further explored.

## Supporting Information

Additional figures, movies, materials and methods

## REFERENCES

1. Laughlin, S. T., Baskin, J. M., Amacher, S. L. & Bertozzi, C. R. In vivo imaging of membrane-associated glycans in developing zebrafish. Science 320, 664–667 (2008).

2. Laughlin, S. T. & Bertozzi, C. R. Imaging the glycome. Proc. Natl. Acad. Sci. U.S.A. 106, 12–17 (2009).

3. Laughlin, S. T. & Bertozzi, C. R. In vivo imaging of Caenorhabditis elegans glycans. ACS Chem. Biol. 4, 1068–1072 (2009).

4. Xie, R. et al. Targeted imaging and proteomic analysis of tumor-associated glycans in living animals. Angew. Chem. Int. Ed. Engl. 53, 14082–14086 (2014).

5. Kang, K. et al. Tissue-based metabolic labeling of polysialic acids in living primary hippocampal neurons. Proc. Natl. Acad. Sci. U.S.A. 112, E241–8 (2015).

6. Prescher, J. A. & Bertozzi, C. R. Chemistry in living systems. Nat. Chem. Biol. 1, 13–21 (2005).

7. Bertozzi, C. R. A decade of bioorthogonal chemistry. Acc. Chem. Res. 44, 651–653 (2011).

8. Capicciotti, C. J. et al. Cell-Surface Glyco-Engineering by Exogenous Enzymatic Transfer Using a Bifunctional CMP-Neu5Ac Derivative. J. Am. Chem. Soc. 139, 13342–13348 (2017).

9. Sun, T. et al. One-Step Selective Exoenzymatic Labeling (SEEL) Strategy for the Biotinylation and Identification of Glycoproteins of Living Cells. J. Am. Chem. Soc. 138, 11575–11582 (2016).

10. Li, J. et al. A Single-Step Chemoenzymatic Reaction for the Construction of Antibody-Cell Conjugates. ACS Cent Sci 4, 1633–1641 (2018).

11. Hong, S. et al. Bacterial glycosyltransferase-mediated cell-surface chemoenzymatic glycan modification. Nat Commun 10, 1799 (2019).

12. Backus, K. M. et al. Uptake of unnatural trehalose analogs as a reporter for Mycobacterium tuberculosis. Nat. Chem. Biol. 7, 228–235 (2011).

13. Tan, H. Y. et al. Direct One-Step Fluorescent Labeling of O-GlcNAc-Modified Proteins in Live Cells Using Metabolic Intermediates. J. Am. Chem. Soc. 140, 15300–15308 (2018).

14. Zon, L. I. & Peterson, R. T. In vivo drug discovery in the zebrafish. Nat Rev Drug Discov 4, 35–44 (2005).

15. Lieschke, G. J. & Currie, P. D. Animal models of human disease: zebrafish swim into view. Nat Rev Genet 8, 353–367 (2007).

16. White, R., Rose, K. & Zon, L. Zebrafish cancer: the state of the art and the path forward. Nat. Rev. Cancer 13, 624–636 (2013).

17. Ramya, T. N. C. et al. In situ trans ligands of CD22 identified by glycan-protein photocross-linking-enabled proteomics. Mol. Cell Proteomics 9, 1339–1351 (2010).

18. Agarwal, P., Beahm, B. J., Shieh, P. & Bertozzi, C. R. Systemic Fluorescence Imaging of Zebrafish Glycans with Bioorthogonal Chemistry. Angew. Chem. Int. Ed. Engl. 54, 11504–11510 (2015).

19. Han, S., Collins, B. E., Bengtson, P. & Paulson, J. C. Homomultimeric complexes of CD22 in B cells revealed by protein-glycan cross-linking. Nat. Chem. Biol. 1, 93–97 (2005).

20. Wang, W. et al. Sulfated ligands for the copper(I)-catalyzed azide-alkyne cycloaddition. Chem Asian J 6, 2796–2802 (2011).

21. North, S. J. et al. Glycomics profiling of Chinese hamster ovary cell glycosylation mutants reveals N-glycans of a novel size and complexity. J. Biol. Chem. 285, 5759–5775 (2010).

22. Petit, D. et al. Integrative view of α2,3-sialyltransferases (ST3Gal) molecular and functional evolution in deuterostomes: significance of lineage-specific losses. Mol. Biol. Evol. 32, 906–927 (2015).

23. Teppa, R. E., Petit, D., Plechakova, O., Cogez, V. & Harduin-Lepers, A. Phylogenetic-Derived Insights into the Evolution of Sialylation in Eukaryotes: Comprehensive Analysis of Vertebrate β-Galactoside α2,3/6-Sialyltransferases (ST3Gal and ST6Gal). Int J Mol Sci 17, (2016).

24. Chang, L.-Y. et al. Novel Zebrafish Mono-α2,8-sialyltransferase (ST8Sia VIII): An Evolutionary Perspective of α2,8-Sialylation. Int J Mol Sci 20, (2019).

25. Kimmel, C. B. & Law, R. D. Cell lineage of zebrafish blastomeres. I. Cleavage pattern and cytoplasmic bridges between cells. Developmental Biology 108, 78–85 (1985).

26. Rillahan, C. D. et al. Global metabolic inhibitors of sialyl- and fucosyltransferases remodel the glycome. Nat. Chem. Biol. 8, 661–668 (2012).

27. Reischauer, S. et al. Cloche is a bHLH-PAS transcription factor that drives haemato-vascular specification. Nature 535, 294–298 (2016).

28. Gore, A. V., Pillay, L. M., Venero Galanternik, M. & Weinstein, B. M. The zebrafish: A fintastic model for hematopoietic development and disease. Wiley Interdiscip Rev Dev Biol 7, e312 (2018).

29. Bertrand, J. Y. et al. Haematopoietic stem cells derive directly from aortic endothelium during development. Nature 464, 108–111 (2010).

30. Foxall, C. et al. The three members of the selectin receptor family recognize a common carbohydrate epitope, the sialyl Lewis(x) oligosaccharide. J. Cell Biol. 117, 895–902 (1992).

31. Springer, T. A. Traffic signals for lymphocyte recirculation and leukocyte emigration: the multistep paradigm. Cell 76, 301–314 (1994).

32. Davidson, A. J. & Zon, L. I. The ‘definitive’ (and ‘primitive’) guide to zebrafish hematopoiesis. Oncogene 23, 7233–7246 (2004).

33. Clements, W. K. & Traver, D. Signalling pathways that control vertebrate haematopoietic stem cell specification. Nat. Rev. Immunol. 13, 336–348 (2013).

34. Baskin, J. M., Dehnert, K. W., Laughlin, S. T., Amacher, S. L. & Bertozzi, C. R. Visualizing enveloping layer glycans during zebrafish early embryogenesis. PNAS 107, 10360–10365 (2010).

35. Dehnert, K. W. et al. Metabolic labeling of fucosylated glycans in developing zebrafish. ACS Chem. Biol. 6, 547–552 (2011).

36. Bardor, M., Nguyen, D. H., Diaz, S. & Varki, A. Mechanism of uptake and incorporation of the non-human sialic acid N-glycolylneuraminic acid into human cells. J. Biol. Chem. 280, 4228–4237 (2005).

37. Gilormini, P.-A. et al. Chemical glycomics enrichment: imaging the recycling of sialic acid in living cells. J. Inherit. Metab. Dis. 41, 515–523 (2018).

38. Shi, S. & Stanley, P. Protein O-fucosyltransferase 1 is an essential component of Notch signaling pathways. PNAS 100, 5234–5239 (2003).

39. Luo, Y., Koles, K., Vorndam, W., Haltiwanger, R. S. & Panin, V. M. Protein O-fucosyltransferase 2 adds O-fucose to thrombospondin type 1 repeats. J. Biol. Chem. 281, 9393–9399 (2006).

40. Wang, X. et al. Dysregulation of TGF-beta1 receptor activation leads to abnormal lung development and emphysema-like phenotype in core fucose-deficient mice. PNAS 102, 15791–15796 (2005).

41. Schneider, M., Al-Shareffi, E. & Haltiwanger, R. S. Biological functions of fucose in mammals. Glycobiology 27, 601–618 (2017).

42. Hellbusch, C. C. et al. Golgi GDP-fucose transporter-deficient mice mimic congenital disorder of glycosylation IIc/leukocyte adhesion deficiency II. J. Biol. Chem. 282, 10762–10772 (2007).

43. Zhou, L. et al. Notch-dependent control of myelopoiesis is regulated by fucosylation. Blood 112, 308–319 (2008).

44. Xia, L., McDaniel, J. M., Yago, T., Doeden, A. & McEver, R. P. Surface fucosylation of human cord blood cells augments binding to P-selectin and E-selectin and enhances engraftment in bone marrow. Blood 104, 3091–3096 (2004).

45. Bigas, A., Robert-Moreno, A. & Espinosa, L. The Notch pathway in the developing hematopoietic system. Int. J. Dev. Biol. 54, 1175–1188 (2010).

46. Myers, J. et al. Fucose-deficient hematopoietic stem cells have decreased self-renewal and aberrant marrow niche occupancy. Transfusion 50, 2660–2669 (2010).

